# A facile method for fluorescent visualization of newly synthesized fibrous collagen by capturing allysine aldehyde groups as cross-link precursors

**DOI:** 10.1101/2025.06.19.660320

**Authors:** Junpei Kuroda, Kazunori K. Fujii, Sugiko Futaki, Azumi Hirata, Yuki Taga, Takaki Koide

## Abstract

The fibrous structures of collagen provide physical strength and stability to tissues and organs. Abnormalities in their orientation, growth, and remodeling cause morphogenetic defects and diseases including fibrosis, highlighting the importance of understanding how collagen fibers are organized within tissues. However, this process remains difficult to study, as methods for fluorescently labeling collagen fibers with simple protocols and visualizing their three-dimensional structure are still limited. Here we describe a convenient method for fluorescent labeling of collagen fibers in vertebrate tissues. Using DAF-FM, a probe originally developed for nitric oxide detection, premature collagen fibers can be visualized through covalent binding to allysine residues, which serve as precursors of collagen cross-linking. We further show that combining two probes with different emission spectra, DAF-FM and DAR-4M, enables pulse–chase labeling of newly synthesized collagen fibers. This approach provides a practical tool for investigating collagen dynamics during tissue development and remodeling.

## INTRODUCTION

The physical properties and proper function of tissues depend heavily on the organization and orientation of extracellular matrix (ECM) components (*1*, *2*), particularly fibrillar collagen. Collagen fibers provide mechanical strength and serve as structural scaffolds that support tissue morphogenesis and homeostasis (*1–6*). Disruptions in collagen fiber formation, alignment, or remodeling contribute to developmental abnormalities and diseases such as osteogenesis imperfecta and fibrosis (*7–10*). Understanding how collagen fibers are organized and remodeled within tissues therefore requires imaging approaches that can visualize their architecture and dynamics *in vivo*.

A variety of methods have been developed to observe collagen fibers in tissues. Immunofluorescent staining remains a widely used approach for visualizing collagen distribution (*11–13*), and recent advances in tissue clearing have improved deep-tissue imaging. Multiphoton microscopy with second-harmonic generation (SHG) detection enables label-free visualization of fibrillar collagen in living tissues and has been instrumental in studying collagen organization across developmental and pathological contexts (*14*, *15*). Genetic approaches using fluorescently tagged collagen have also provided valuable insights into collagen dynamics *in vivo* (*16–19*). In addition, low–molecular-weight chemical approaches have recently emerged, including staining with the synthetic colorant Fast Green FCF, which enables deep-tissue visualization of collagen networks but requires multiple preparation steps and loses specificity under hydrophilic conditions (*20*). Other chemical strategies targeting precursors of collagen cross-linking have also been reported (*21–25*), yet these methods still do not provide a rapid and broadly applicable means for fluorescent labeling of collagen fibers across diverse vertebrate tissues. In particular, techniques that allow selective visualization of newly synthesized collagen fibers and enable pulse–chase analysis remain limited.

Diaminofluorescein-FM (DAF-FM), a probe originally developed for nitric oxide (NO) detection (*26–28*), has been reported to fluorescently label collagen-rich structures such as notochord, cartilage, bone, and actinotrichia in zebrafish (*29–33*). More recently, DAF-FM was shown to label collagen fibers in axolotl skin (*34*). These observations suggested that DAF-FM may interact with collagen fibers independently of NO, raising the possibility that this probe could be repurposed as a chemical tool for collagen imaging. However, although previous studies reported DAF-FM staining of collagen-rich structures, the underlying chemical mechanism and its utility for time-resolved labeling of collagen fibers have not been systematically examined.

In this study, we investigated the interaction between DAF-FM and collagen fibers and evaluated its utility as a simple method for fluorescent labeling of collagen architecture. We found that DAF-FM labels collagen fibers through covalent binding to allysine residues, which serve as precursors of collagen cross-linking. This reaction enables rapid staining of collagen fibers in cultured fibroblasts while maintaining cell viability. We further demonstrate that combining DAF-FM with the related probe diaminorhodamine-4M (DAR-4M) allows pulse–chase labeling of newly synthesized collagen fibers, enabling temporal analysis of collagen fiber growth *in vitro*.

We also show that DAF-FM DA, a membrane-permeable derivative, enables clear three-dimensional visualization of collagen fibers deep within mouse tissues. In addition, DAF-FM DA labels collagen fibers in other vertebrates, including zebrafish and axolotl, demonstrating its broad applicability. Finally, we apply the dual-probe pulse–chase strategy to *in vivo* imaging of collagen fibers in zebrafish bone, allowing temporal visualization of collagen deposition during tissue growth.

Together, these findings establish DAF-FM–based labeling as a practical approach for visualizing collagen fibers and tracking their dynamics across a range of vertebrate tissues. This method complements existing imaging strategies and provides a simple chemical tool for studying collagen organization and remodeling during development and tissue homeostasis.

## RESULTS

### DAF-FM and DAR-4M enable fluorescent labeling of collagen fibers formed by cultured cells

Previous studies have reported that DAF-FM DA, a known NO detection probe, is effective for fluorescent staining of collagen fiber-rich tissues such as the notochord, cartilage, bone, and skin (*29–34*). Based on the results of these studies, we hypothesized that this probe could fluorescently stain collagen fibers independent of NO. In this study, we first tested this possibility by culturing mouse fibroblasts with high collagen fiber production activity. In this experiment, we used DAF-FM, a membrane-impermeable version of the DAF-FM DA. Before starting the culture experiment, we coated the culture dishes with purified collagen to determine whether the substrates would be fluorescently stained with DAF-FM. We confirmed that DAF-FM exhibited minimal reactivity with the purified collagen used as the culture substrate (Fig. S1). Therefore, we seeded mouse embryonic fibroblasts (MEFs) on gelatin-coated culture dishes and stained them with DAF-FM, while the cells were alive after culture (Fig. 1A). The network of fiber structures formed by MEFs was fluorescently labeled by DAF-FM (Fig. 1B). To determine whether these fibrous structures were collagen fibers, we conducted double staining with DAF-FM and Bin*d*COL, a cyclic peptide that hybridizes to the denatured portions of collagen (*35*). Importantly, fibers labeled with DAF-FM were simultaneously labeled with Bin*d*COL after denaturation by heat treatment, indicating that DAF-FM fluorescently stained collagen fibers formed by cultured fibroblasts (Fig. 1B). We also found that when co-stained with DAF-FM and its derivative DAR-4M, the two probes fluorescently labeled the same fiber structures (Fig. S2). We also performed double staining with DAR-4M using an antibody against type I collagen (Col1) and found that DAR-4M staining merged very well with Col1 antibody staining (Fig. S2). These results indicated that DAF-FM and DAR-4M can fluorescently label collagen fibers produced by fibroblasts under culture conditions using a simple method.

**Figure 1.**
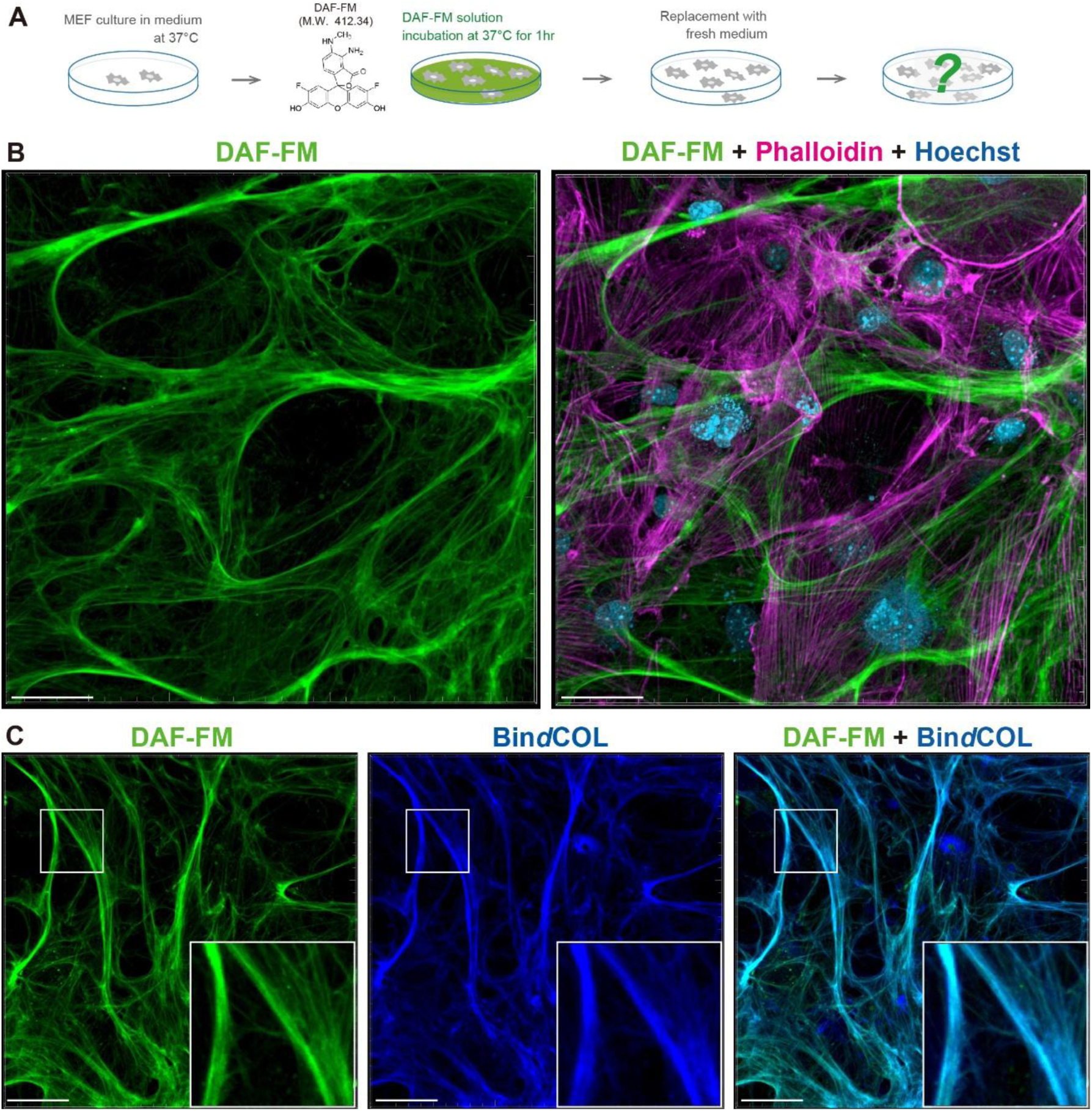
A rapid and simple method for the visualization of the collagen fibers formed by culture cells using DAF-FM. (A) Schematic diagram of DAF-FM staining for the collagen fibers formed by MEFs. Fully proliferated MEFs after 1–2 weeks of culture were incubated in 10 μM DAF-FM solution for 1 h. After the incubation, DAF-FM solution was replaced with fresh medium and the collagen fibers visualized by DAF-FM were observed. (B) Representative fluorescent images of the collagen fibers labeled by DAF-FM (green) at culture day 10. Actin cytoskeleton was stained with phalloidin (magenta), and nucleus was stained with Hoechst (blue). (C) Representative fluorescent images of the collagen fibers co-stained with DAF-FM (green) and Bin*d*COL (blue). Scale bar = 50 μm.

Next, we investigated whether the fluorescence of collagen fibers observed after DAF-FM staining was due to NO produced by the cells. Therefore, to clarify the involvement of NO, we treated MEFs under culture conditions to remove NO and observed the fluorescence of the fibers after DAF-FM staining. No significant difference was observed in the intensity of the fluorescent signal from the fibers under NO removal conditions compared with the control (Fig. S3), indicating that NO is not involved in the DAF-FM-induced fluorescence emission of collagen fibers.

We also investigated the effects of DAF-FM staining on cell activity. We treated cultured MEFs with DAF-FM and performed a BrdU incorporation assay to assess cell proliferation (Fig. S4). There was no significant difference in BrdU incorporation between cells treated with DAF-FM and untreated cells (Fig. S4). To determine the effect of DAF-FM on cell survival, we examined whether apoptosis was enhanced by DAF-FM treatment. We used nuclear staining reagents to detect apoptosis and found no significant difference in cell death between cells treated with DAF-FM and untreated cells (Fig. S4). Based on these results, we conclude that DAF-FM does not adversely affect cell activity, at least at low concentrations, and that this probe allows the visualization of collagen fibers produced by living cells independent of NO.

### Fluorescent labeling of collagen fibers with DAF-FM via interaction with cross-linking intermediate structures

To clarify the affinity and specificity of DAF-FM for collagen fibers, we attempted to identify the mechanism by which this fluorescent probe labeled collagen fibers. DAF-FM is widely used as a probe that reacts with NO to emit fluorescence (*36*), but the result in Fig. S3 indicated that NO was not involved in the fluorescent labeling of collagen fibers by DAF-FM. DAF-FM was recently reported to emit green fluorescence by binding to aldehydes and NO (*37*). In addition, it is known that ε-amino groups of lysine residues in the N- and C-terminal non-triple helical domains (telopeptides) of collagen α-chain are modified to aldehyde groups by lysyl oxidase (LOX), and collagen molecules undergo intra- and intermolecular cross-linking via these aldehyde groups, resulting in the polymerization and growth of collagen fibers (*38–40*). These observations suggest that DAF-FM fluorescently labels collagen fibers by interacting with aldehydes, which are intermediate structures in collagen cross-links. Therefore, we examined the fluorescence after DAF-FM staining under conditions that inhibited collagen cross-linking. We treated MEFs in culture condition with ꞵ-aminopropionitrile (BAPN), known as an inhibitor of LOX (*41*), in this experiment (Fig. 2A). The inhibition of collagen cross-linking by treatment with BAPN suppressed the fluorescent labeling of fibers by DAF-FM (Fig. 2B). However, when MEFs were cultured under normal conditions and stained with DAF-FM in the presence of BAPN, a clear fluorescence of collagen fibers was observed, similar to the control (Fig. S5A). These results suggested that the fluorescent labeling of collagen fibers by DAF-FM was closely related to the modification of collagen by LOX.

**Figure 2.**
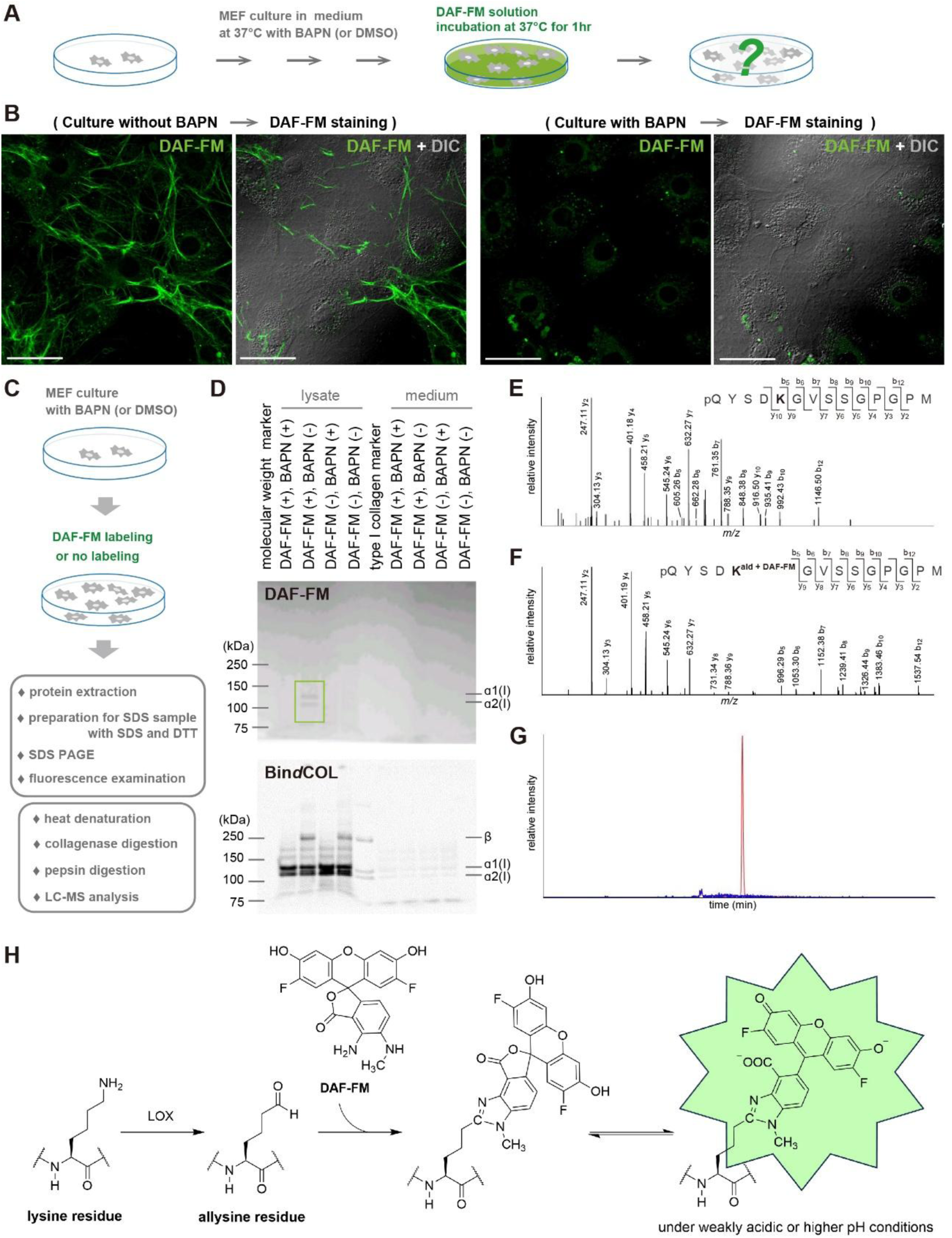
Mechanism of DAF-FM fluorescence targeting collagen cross-linking. (A) Schematic diagram of DAF-FM staining for the collagen fibers formed by MEFs under the culture condition with BAPN. (B) Representative fluorescent images of the collagen fibers labeled by DAF-FM (green) under the culture condition without BAPN and with BAPN. Scale bar = 50 μm. (C) Experimental workflow for sample preparation for SDS-PAGE and LC-MS analysis. (D) SDS-PAGE analysis of DAF-FM-labeled collagen. DAF-FM fluorescence of proteins derived from the lysate and medium at each condition was examined (upper panel), and Bin*d*COL staining of the same protein samples at each condition was performed (lower panel). (E) MS/MS spectra of α2-N telopeptide containing lysine (*m*/*z* 696.8085, *z* = 2) derived from control (non-labeled) samples. (F) MS/MS spectra of α2-N telopeptide containing DAF-FM-labeled allysine (*m*/*z* 892.3239, *z* = 2) derived from DAF-FM-labeled samples. (G) Monoisotopic extracted ion chromatograms of the DAF-FM-labeled α2-N telopeptide (*m*/*z* 892.3352 ± 0.02, *z* = 2) for the control (blue) and DAF-FM-labeled sample (red). (H) Proposed mechanism of collagen fiber visualization with DAF-FM.

Next, we attempted to determine the specific target sites of the collagen cross-linking structure with which DAF-FM interacts. First, proteins were extracted from MEF culture dishes treated with DAF-FM and SDS-PAGE was performed under reducing conditions using DTT to examine whether DAF-FM fluorescence was retained (Fig. 2C). As a result, when MEFs were cultured in the absence of BAPN and stained with DAF-FM, specific bands were detected in the lysate-derived sample at the positions of molecular weights predicted to be α1(ǀ) and α2(ǀ) chains (Fig. 2D). However, these specific bands were not detected in lysate-derived samples when the same experiment was performed in the presence of BAPN (Fig. 2D, fig. S5B). We tested whether the same bands could be detected using Bin*d*COL after the same experiment. In the lysate-derived sample, specific bands labeled with Bin*d*COL were detected at the positions of molecular weights predicted to be α1(ǀ) and α2(ǀ) (Fig. 2D, Fig. S5B). These results indicate that under culture conditions in the presence of BAPN, the production of collagen fibers occurs normally, but the inhibition of cross-linking suppresses the fluorescence caused by DAF-FM. Furthermore, the fluorescence caused by DAF-FM in the two specific bands did not disappear after pepsin treatment, indicating that DAF-FM specifically reacted with protease-resistant collagen molecules because of their triple helical structure (Fig. S5C). In addition, our results suggest that DAF-FM fluoresces through covalent binding to specific structures that occur during the cross-linking formation process.

We further attempted to identify the binding site of DAF-FM by LC-MS analysis (Fig. 2C). In this experiment, we analyzed three collagenase/pepsin-digested peptides derived from telopeptide domains of type I collagen (α1-N telopeptide, α1-C telopeptide and α2-N telopeptide) which contain lysine or hydroxylysine participating in cross-link formation. Telopeptidyl lysine residues are not hydroxylated in the skin (*42*), but are largely converted to hydroxylysine in the bone (*43*). Thus, we analyzed lysine-containing telopeptides in the MEF samples. MS/MS sequence analysis confirmed that DAF-FM binds to the aldehyde groups of lysine residues within the α2-N telopeptide in the DAF-FM-labeled sample (Fig. 2E and F). The extracted ion chromatograms showed that a strong peak for this telopeptide labeled with DAF-FM was detected only in the DAF-FM-labeled sample (Fig. 2G). Extracted ion chromatogram peaks corresponding to the theoretical molecular weight of α1-N and α1-C telopeptides attached with DAF-FM were only slightly detected in the DAF-FM-labeled sample, and MS/MS sequence confirmation could not be performed (data not shown). The mass shift observed in the MS/MS spectra suggested that DAF-FM reacted with aldehyde groups to form fluorophores via a previously reported mechanism (*37*). The predicted fluorophore was generated by reacting DAF-FM with the aldehyde group of the allysine residue (Fig. 2H). Based on these results, we concluded that DAF-FM fluorescently labeled collagen fibers by covalently binding to the aldehyde groups of allysine (and possibly hydroxylysine) residues in telopeptides, which are intermediate structures in collagen cross-linking.

### Fluorescent visualization of collagen fibers with DAF-FM DA in cartilage and notochord of mouse embryos

Using cultured mouse cells, we successfully demonstrated that DAF-FM fluorescently labels collagen fibers through a mechanism distinct from that of NO detection. We then attempted DAF-FM staining of mouse tissues to determine whether this probe is also useful for the fluorescent imaging of collagen fibers *in vivo*. In previous studies, DAF-FM DA, which exhibits plasma membrane permeability (*27*), has been used for whole-body fluorescence staining of zebrafish (*29–32*). Therefore, we assumed that DAF-FM DA has high tissue penetration and decided to use this probe for the fluorescent staining of mouse tissues.

We first examined whether two collagen-rich tissues, the cartilage and notochord, could be fluorescently stained with DAF-FM DA in mouse embryos. Unfixed embryos at embryonic day (E) 14.5 were incubated overnight in diluted DAF-FM DA solution (Fig. 3A and B). After incubation, the tails of the embryos were removed to examine the three-dimensional orientation of the collagen fibers in the cartilage and notochord. Confocal microscopy revealed strong green fluorescent signals in the cartilage primordium and notochord (Fig. 3C-C”, Movie S1). To determine whether the DAF-FM DA signals were derived from collagen fibers, sections of DAF-FM DA-labeled embryos were incubated with an anti-type II collagen (Col2) antibody. The reactivity of type II collagen, the major extracellular matrix component of cartilage, was detected in the area where DAF-FM DA fluorescence was observed (Fig. 3D), indicating that DAF-FM DA also fluorescently labeled developing collagen fibers in mouse embryos.

**Figure 3.**
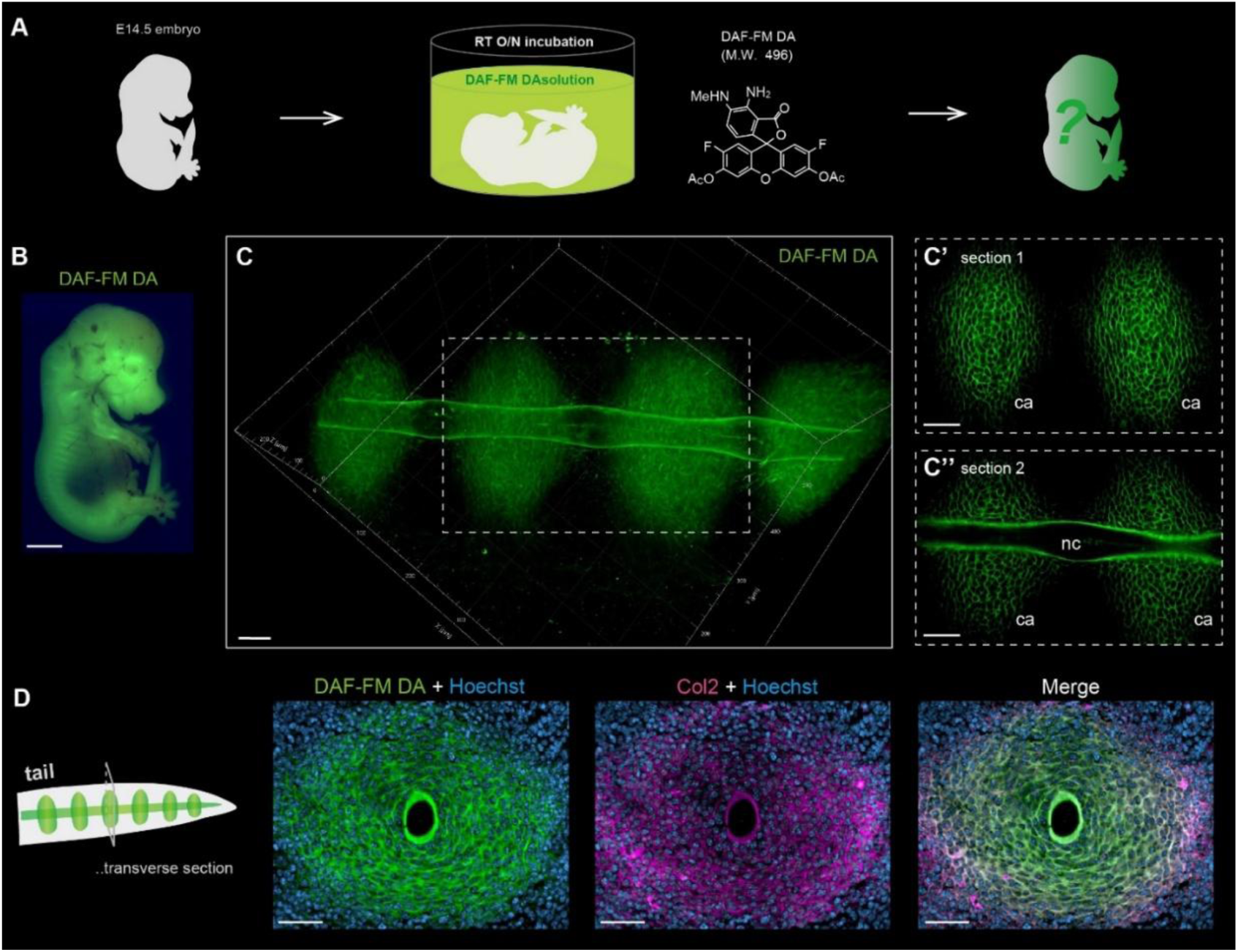
DAF-FM DA enables clear fluorescent visualization of the collagen fibers in cartilage and notochord of mouse embryos with a simple method. (A) Schematic diagram of DAF-FM DA staining for the collagen fibers in mouse embryos. (B) The fluorescent image of the E14.5 embryo after DAF-FM DA staining. (C) The 3D fluorescent image of the embryonic tail stained with DAF-FM DA. Slice images of the area within the white dotted box are shown in (C’ and C’’). ca, cartilage; nc, notochord. (D) The fluorescent images of antibody staining of the cryo-section samples of the tail. The tissue sections labeled with DAF-FM DA (green) were stained with anti-Col2 antibody (magenta) and Hoechst (blue). Scale bar = 2 mm (B) and 50 μm (C to D).

### Fluorescent visualization of collagen fibers with DAF-FM DA in tendons and cartilage of postnatal mouse

Next, we investigated whether DAF-FM DA is useful for fluorescent visualization of collagen fibers in postnatal mice. To simultaneously observe cartilage and other collagen-rich tissues, such as tendons, the tails of newborn mice were stained with DAF-FM DA. After incubation in the DAF-FM DA solution, the tail samples were cleared with transparency reagents. Strong DAF-FM DA signals were observed in the tendons and fibrocartilage of intervertebral discs deep inside the tail (Fig. 4A). To simultaneously observe the distribution of collagen fibers and surrounding cells, nuclear staining was performed on tail samples labeled with DAF-FM DA (Fig. 4B). Each XY cross-sectional image scanned using confocal microscopy displayed clearly labeled fiber structures in the tendon and intervertebral fibrocartilage (Fig. 4C, Movie S2). The nuclei were positioned in the gaps between the fibers (Fig. 4C). We further confirmed that the fluorescence of DAF-FM DA overlapped well with the SHG signal in tissue sections of the intervertebral discs (Fig. 4D). The orientation patterns of fibers labeled with each fluorescent dye were consistent (Fig. 4D). The SHG signal intensity was not uniform and differed for each fiber, whereas the DAF-FM DA signal was detected uniformly throughout the fiber (Fig. 4D). In conclusion, we demonstrated that DAF-FM DA can be used to observe collagen fibers in postnatal mouse tissues with high resolution and identify individual fibers using an easy method.

**Figure 4.**
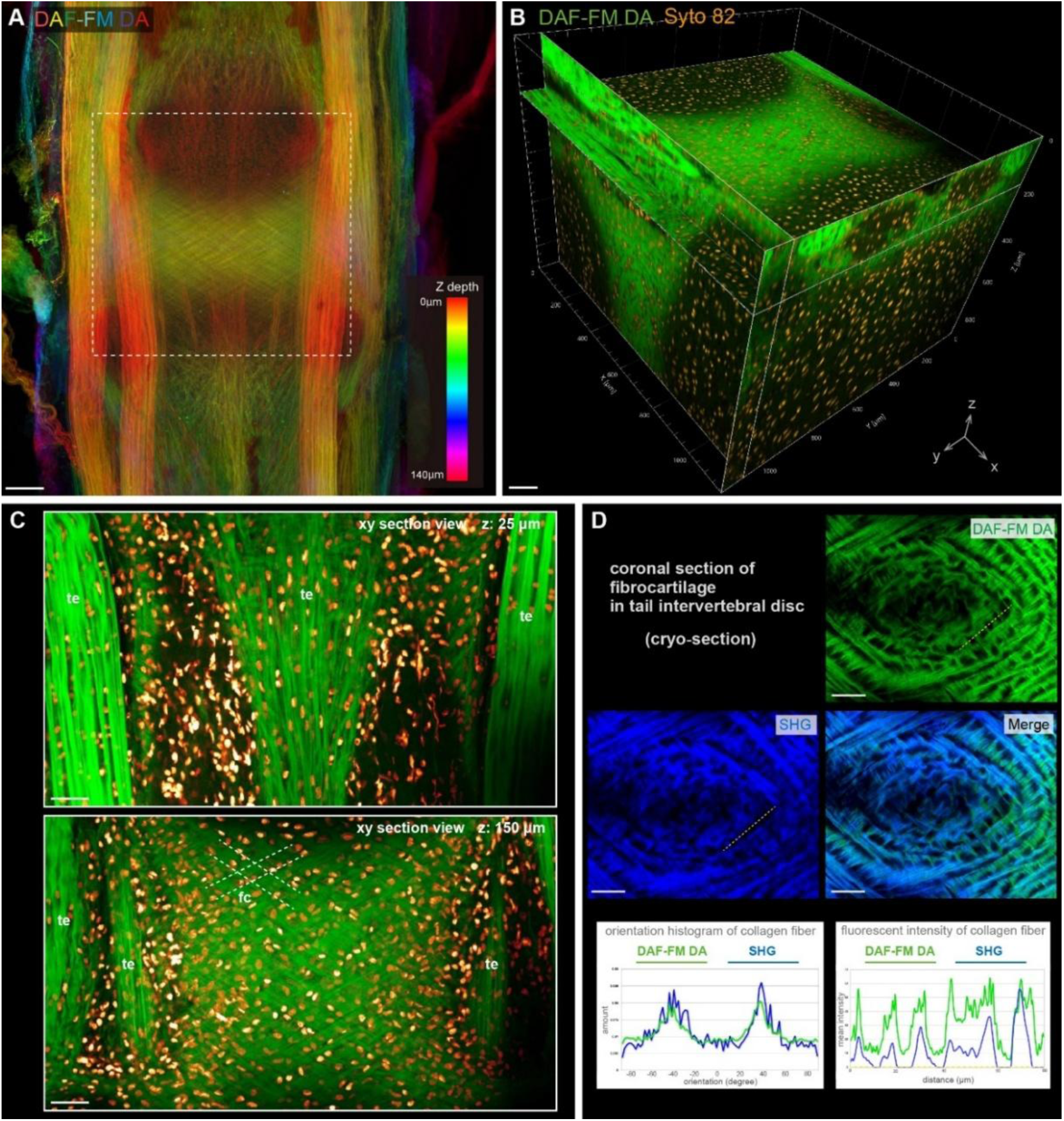
Fluorescent visualization of the collagen fibers in tendon and cartilage of postnatal mouse using DAF-FM DA. (A) The confocal fluorescent image with depth color-coded MIP of the tendon and cartilage in postnatal mouse tail stained with DAF-FM DA. (B) The 3D reconstructed confocal image in the area within the white dotted box of (A). The collagen fibers were stained with DAF-FM DA (green), and all nuclei were stained with Syto 82 (orange). (C) XY sectional views at an intervertebral region. te, tendon; fc, fibrocartilage. White dotted lines indicate the fibrocartilage orientation. (D) The fluorescent images of cryo-section samples at the tail intervertebral disc region. Orientation and fluorescent intensity of the collagen fibers visualized with DAF-FM DA (green) and SHG (blue) were plotted, respectively. Scale bar = 100 μm (A to C) and 40 μm (D).

### Fluorescent visualization of collagen fibers in aquatic vertebrates using DAF-FM DA

To further demonstrate the utility of collagen fiber imaging using DAF-FM DA, we performed DAF-FM DA staining on the tissues of other vertebrates. Recently, Kuroda et al. reported that DAF-FM DA labels actinotrichia oriented at the tips of zebrafish fins (*32*). More recently, Ohashi et al. reported that this probe fluorescently labels the collagen fiber network that develops three-dimensionally in the dermis of the axolotl (*34*). We investigated whether the probe labeled collagen fibers in other collagen-rich tissues by staining the entire body of these animals. In this experiment, live juvenile zebrafish and axolotl were incubated in DAF-FM DA solution (Fig. S6A and B). First, we confirmed that DAF-FM DA fluorescently visualized the lattice-like collagen fiber structures that developed in the dermis of juvenile zebrafish and axolotls, consistent with a recent report (Fig. S6C and D). Additionally, tendons at the myoseptal junction were visualized using strong fluorescence (Fig. S6C). In addition, clear fluorescence was observed in the tendons that developed in the joint regions of each fin bone in the zebrafish (Fig. S6C). Furthermore, the tendons and ligaments in the digits of the axolotl were visualized using DAF-FM DA with distinct signals (Fig. S6F). These results demonstrate that DAF-FM DA can be used for whole-body staining of collagen fibers in the tissues of living aquatic vertebrates.

### Pulse-chase observation using two-color fluorescent probes to analyze the growth dynamics of collagen fiber *in vitro* and *in vivo*

Finally, we performed pulse-chase observation, which tracks the growth process of collagen fibers, using two fluorescent probes of different colors: DAF-FM and DAR-4M. For this experiment, it was essential that the fluorescence of the labeled fibers remain stable over an extended period without fading. Therefore, we fluorescently labeled collagen fibers with DAF-FM only once during MEF culture and examined whether the fluorescence faded by temporal observations in the same area (Fig. S7A). The fluorescence of the collagen fibers labeled with DAF-FM barely faded during the culture process after staining (Fig. S7B). We then attempted to color-code collagen fibers during their growth using a combination of the two fluorescent probes at different times during the culture process (Fig. 5A). We successfully distinguished between old fibers, formed four days ago) and newly formed fibers by performing DAF-FM and DAR-4M staining at different time points (Fig. 5B). Our results demonstrated the growth of a network of collagen fibers through the process of thickening by the deposition of new collagen around old fibers (Fig. 5B’ asterisk) and the formation of new fibers near old fibers (Fig. 5B’ arrowheads).

**Figure 5.**
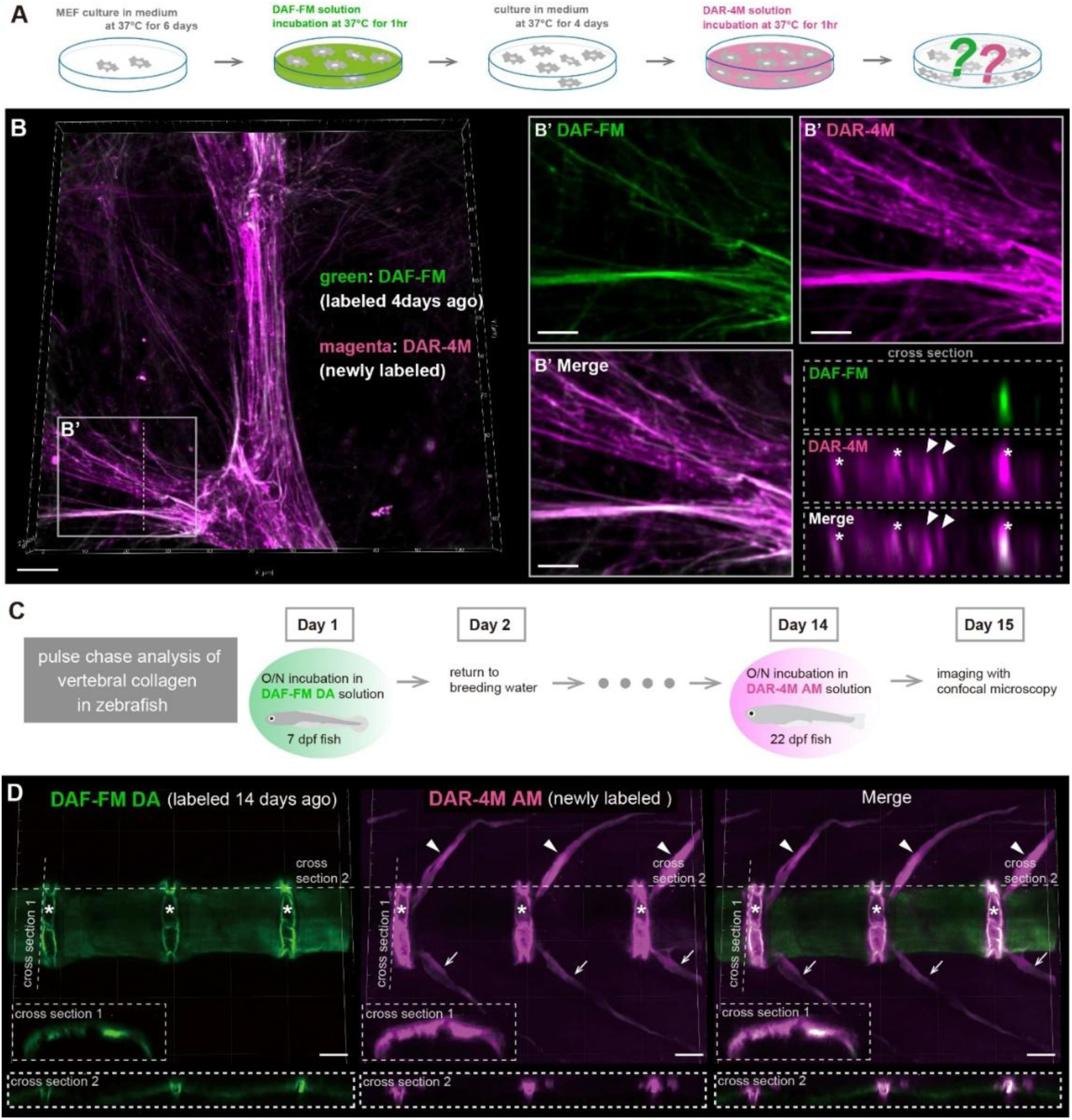
Pulse-chase observation using two different fluorescent probes to understand growth manner of collagen fibers *in vitro* and *in vivo*. (A) Schematic diagram illustrating the pulse-chase observation of the collagen fibers formed by MEFs, using DAF-FM and DAR-4M. MEF cultured for 6 days were first stained with DAF-FM. After DAF-FM staining, the staining solution was replaced with fresh medium and MEFs were incubated for 4 days. Subsequently, DAR-4M staining was performed, and the fluorescence of collagen fibers was observed. (B) Representative fluorescent images of the collagen fibers labeled by DAF-FM (green, labeled 4 days ago) and DAR-4M (magenta, newly labeled). Magnified images in the white box area and cross section images at the position of the white dotted line are shown in the right panels. Asterisks indicate the fibers that have thickened due to the additional growth of older fibers, and arrowheads indicate newly formed fibers between old fibers. (C) Schematic diagram illustrating the pulse-chase observation of the collagen fibers during the formation of vertebral bones in zebrafish, using DAF-FM DA and DAR-4M AM. Living zebrafish larvae at 7 dpf were first stained with DAF-FM DA. After the staining, the fish were returned to fresh tank water and bred for 2 weeks. Subsequently, DAR-4M AM staining was performed, and the fluorescence of the collagen fibers around the vertebrae was observed. (D) Representative fluorescent images of the collagen fibers around the vertebrae labeled by DAF-FM DA (green, labeled 14 days ago) and DAR-4M AM (magenta, newly labeled). Cross section images at the position of the white dotted lines in each fluorescent image are shown in the lower panels. Asterisks indicate the intervertebral regions. Arrowheads and arrows indicate neural spines and hemal spines, respectively. Scale bar = 10 μm (B), 5 μm (B’) and 50 μm (D).

We also attempted to perform pulse-chase observations of collagen fibers in living animal tissues. In the present study, we focused on vertebral development in zebrafish. We first evaluated the reactivity of the two probes to the collagen fibers distributed in the notochord 7 days post-fertilization (dpf) before the start of vertebral calcification (Fig. S8A). At 7 dpf, strong fluorescence was detected in the notochord following DAF-FM DA staining, whereas only weak fluorescence was detected following DAR-4M AM (acetoxymethyl ester) staining (Fig. S8A). In contrast, at the advanced calcification stage around the notochord associated with vertebral bone formation, strong fluorescence was detected in the intervertebral regions and neural and hemal spines after staining with both probes (Fig. S8B). However, at this stage, no clear fluorescence was observed in the central region when either probe was used (Fig. S8B). We found that the fluorescence signals in the intervertebral regions and spines emitted with DAF-FM DA were similarly detected with collagen hybridizing peptide (CHP), a peptide that binds to denatured collagen chains (*44*) and staining after denaturation by heat treatment (Fig. S8C). To clarify the dynamics of the distribution and production of collagen fibers during early osteogenesis in the vertebral region, we first incubated zebrafish larvae at 7 dpf in a DAF-FM DA solution and labeled the collagen fibers distributed in the notochord with green fluorescence (Fig. 5C). After staining with DAF-FM DA, the fish were bred in a circulating system tank for 2 weeks. Subsequently, the fish were incubated in the DAR-4M AM solution (Fig. 5C). Collagen fibers distributed at the notochord, initially labeled with DAF-FM DA, exhibited a remarkable change in their distribution pattern during the 2 weeks of growth (Fig. 5D). The fluorescence of the collagen fibers uniformly labeled at the notochord before bone formation emitted a strong signal, specifically in the intervertebral disc region, during osteogenesis (Fig. 5D). This indicates that collagen fibers uniformly distributed around the notochord at the larval stage were reorganized during bone formation and reused as structurally specific components of the intervertebral disc. In contrast, the collagen fibers labeled with DAR-4M AM showed more extensive fluorescence in the intervertebral region compared to those with DAF-FM DA (Fig. 5D). Furthermore, no obvious signal was observed in the central region, whereas strong fluorescence labeled with DAR-4M AM was observed in the neural and hemal spines (Fig. 5D). This result suggests that in the early stages of vertebral formation, collagen production is more active in the spinal and intervertebral regions than in the vertebral bodies. In summary, we successfully tracked the changes in collagen distribution patterns and visualized the active areas of collagen production in living zebrafish by applying the two probes at different times during osteogenesis. Hence, we demonstrated that DAF-FM DA and DAR-4M AM are useful for the pulse-chase observation of collagen fibers during the growth of living tissues.

**Fig. S1.**
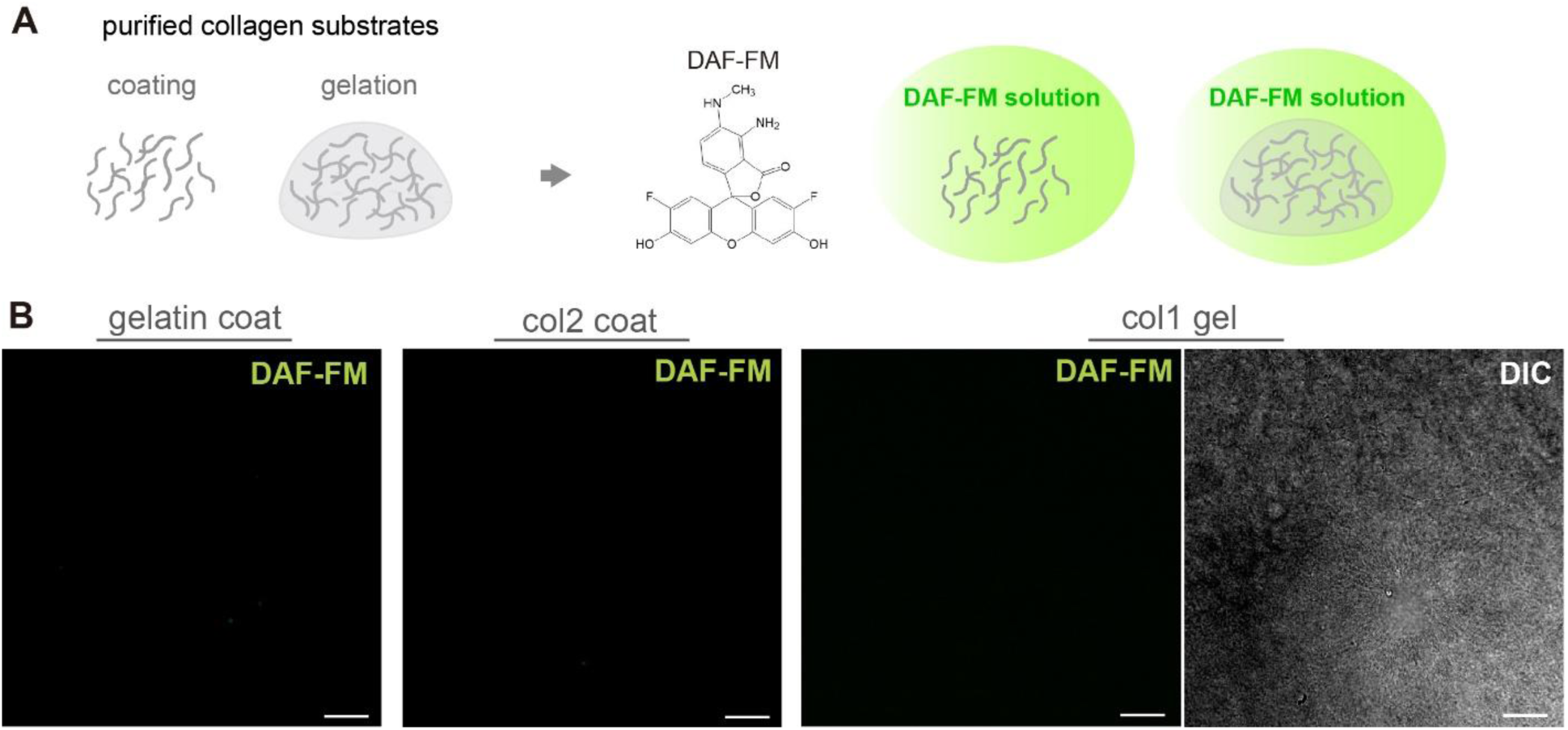
Reactivity of DAF-FM with purified collagen. (**A**) Schematic diagram of DAF-FM staining for the purified collagen substrates. (**B**) Representative confocal images of the purified collagen after DAF-FM staining. Scale bar = 50 μm.

**Fig. S2.**
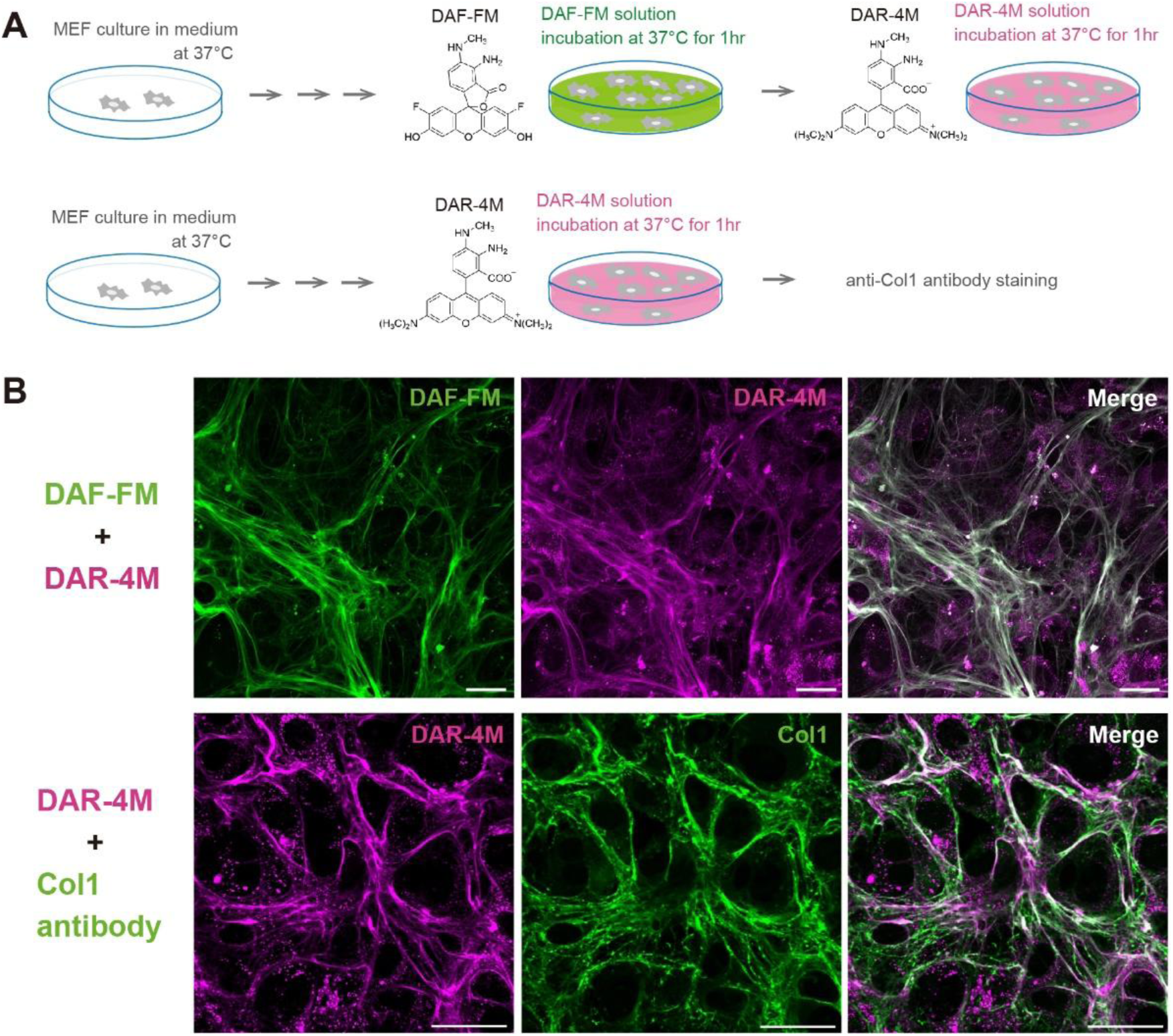
Fluorescent staining of the collagen fibers formed by culture cells using DAR-4M. (**A**) Schematic diagram of DAR-4M staining for the collagen fibers formed by MEFs. (**B**) Upper panels show representative fluorescent images of the collagen fibers visualized with DAF-FM (green) and DAR-4M (magenta) at culture day 10. Both probes visualized the same fibers. Lower panels show representative fluorescent images of the collagen fibers visualized with DAR-4M (magenta) and anti-Col1 antibody staining at culture day 10. The fluorescent signals of DAR-4M and anti-Col1 antibody staining were merged well on the same fibers. Scale bar = 50 μm.

**Fig. S3.**
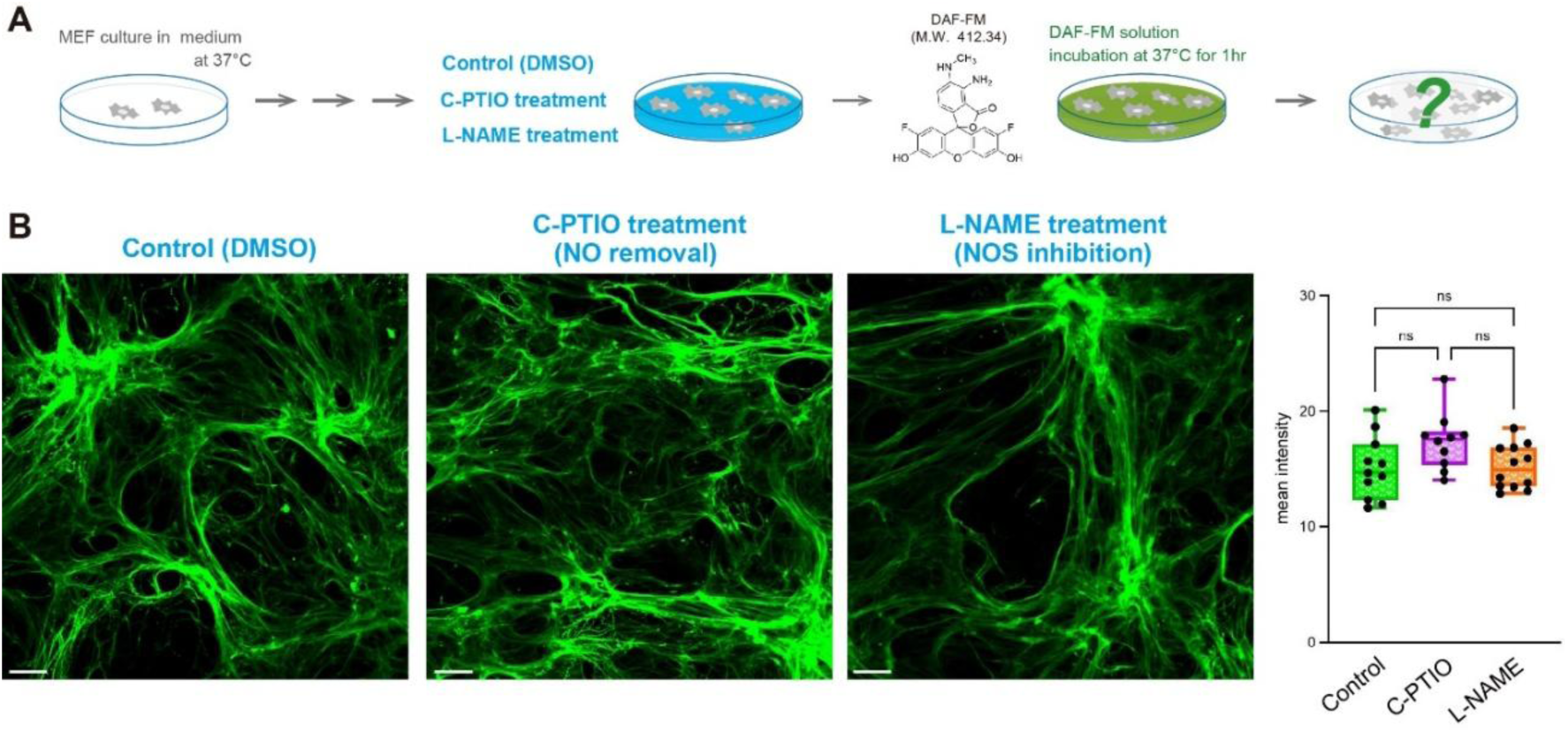
NO-independent fluorescent visualization of collagen fibers by DAF-FM. (**A**) Schematic diagram of DAF-FM staining under NO removal conditions for the collagen fibers formed by MEFs. (**B**) Representative fluorescent images of the collagen fibers visualized with DAF-FM (green) at culture day 10. Clear fluorescent signals of collagen fibers stained with DAF-FM were detected under C-PTIO (NO removal) and L-NAME (NOS inhibition) treatment conditions, similar to the control. The fluorescence intensity plots for each condition are shown in the right panel. Scale bar = 50 μm.

**Fig. S4.**
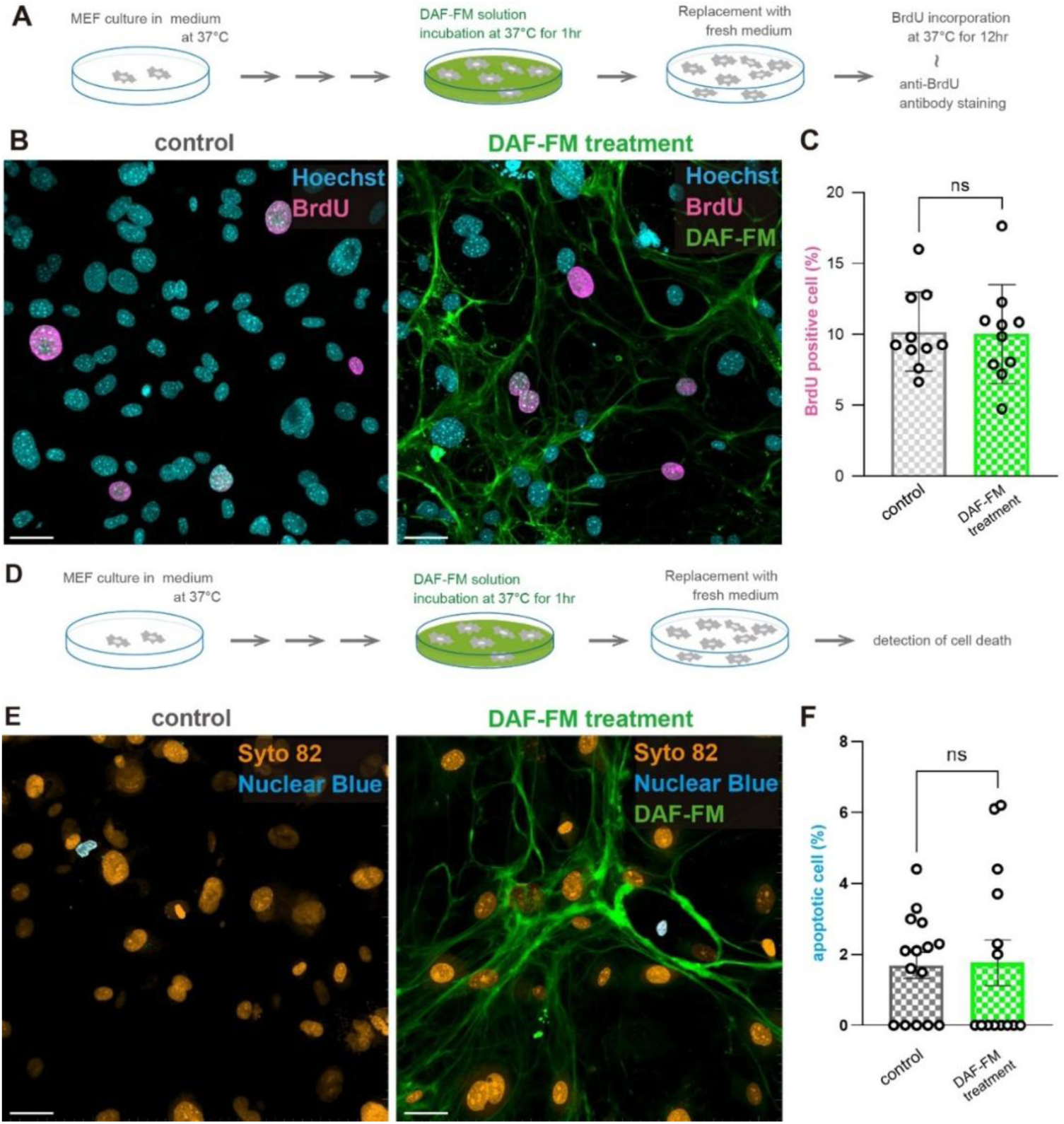
No negative effects on cell activities with DAF-FM treatment. (**A**) Experimental workflow to investigate the effect of DAF-FM treatment on cell division. (**B**) Representative images of anti-BrdU antibody staining. Cultured MEFs were incubated with DMSO (control) or DAF-FM solution. All cell nuclei were stained with Hoechst (blue), and cell nuclei incorporating BrdU were stained with anti-BrdU antibody (magenta). Collagen fibers were stained with DAF-FM (green). (**C**) Number of BrdU-positive cells was counted under each condition. There was no significant difference in the percentage of BrdU-positive cells between the conditions. (**D**) Experimental workflow to investigate the effect of DAF-FM treatment on cell death. (**E**) Representative images of Nuclear Blue staining. Cultured MEFs were incubated with DMSO (control) or DAF-FM solution. All cell nuclei were stained with Syto 82 (orange), and cell nuclei of apoptotic cells were stained with Nuclear Blue (blue). Collagen fibers were stained with DAF-FM (green). (**F**) Number of apoptotic cells was counted under each condition. There was no significant difference in the percentage of apoptotic cells between the conditions. Scale bar = 50 μm.

**Fig. S5.**
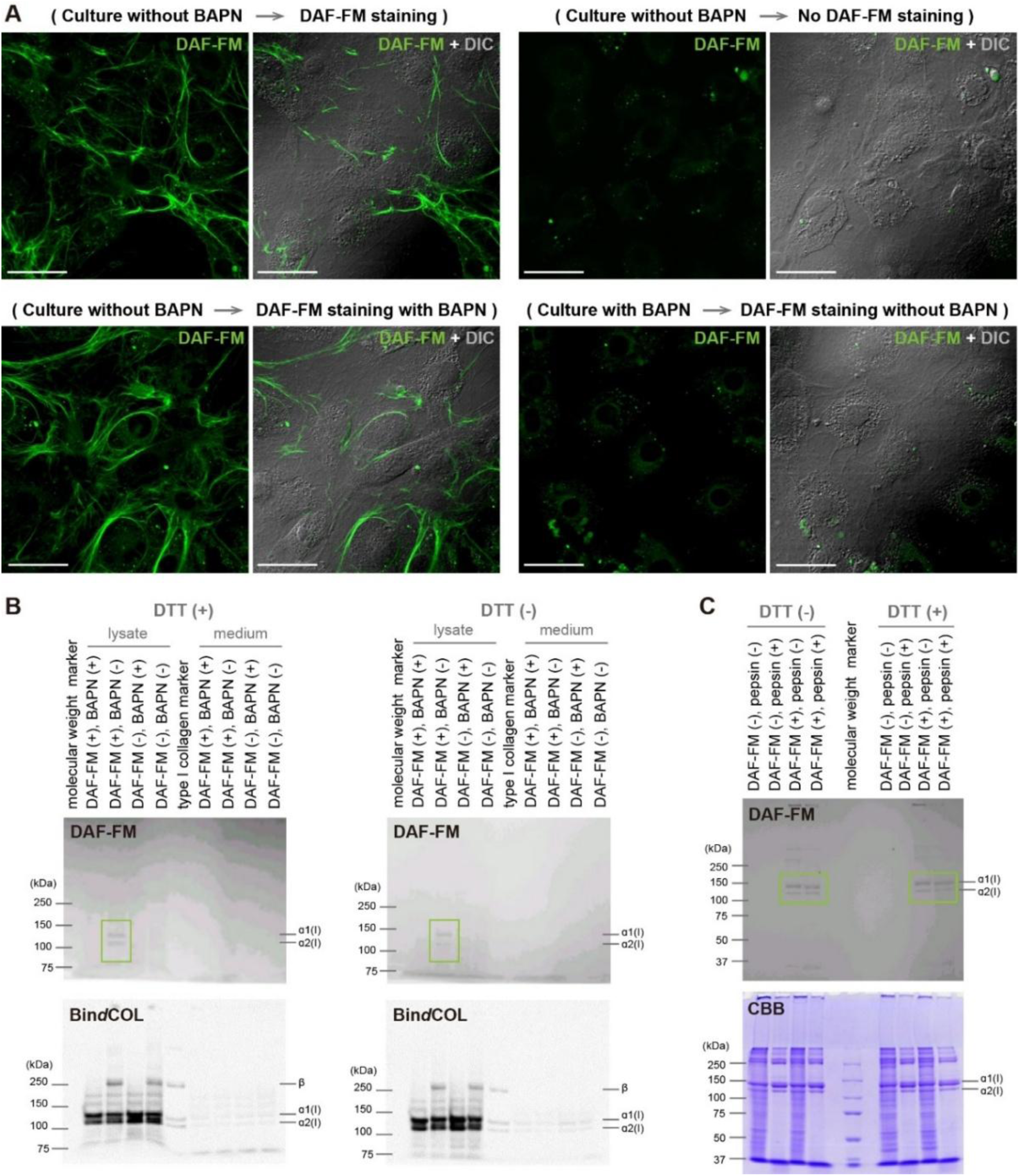
DAF-FM fluorescence suppressed by inhibition of collagen cross-linking formation. (**A**) Representative fluorescent images of the collagen fibers labeled by DAF-FM (green) under the various culture conditions. DAF-FM fluorescence of the collagen fibers produced by MEFs is not suppressed when BAPN is present only during DAF-FM staining, but it is significantly suppressed when BAPN is present during MEF culture. Scale bar = 50 μm. (**B**) SDS-PAGE analysis of DAF-FM-labeled collagen with or without DTT. DAF-FM fluorescence of proteins produced by MEFs in the presence or absence of BAPN was examined (upper panel), and Bin*d*COL staining of the same protein samples was performed (lower panel). (**C**) DAF-FM fluorescence of pepsin-digested or undigested proteins produced by MEFs was examined (upper panel), and CBB staining of the same protein samples was performed (lower panel).

**Fig. S6.**
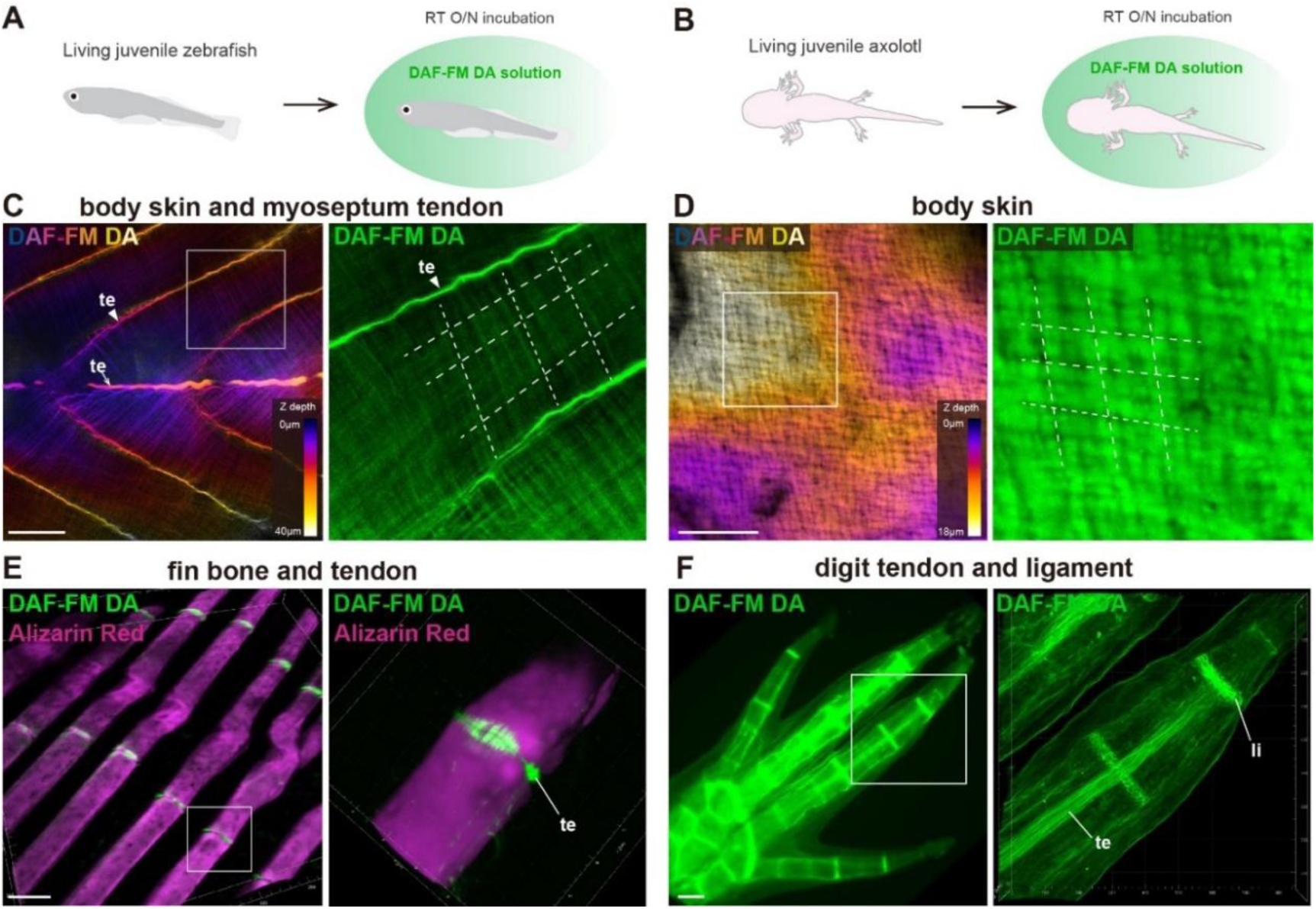
Fluorescent visualization of collagen fibers in zebrafish and axolotl using DAF-FM DA. (**A, B**) Schematic diagram of DAF-FM DA staining for the collagen fibers in zebrafish and axolotl. Living juvenile zebrafish and axolotl were incubated overnight in 10μM DAF-FM solution. After the incubation, DAF-FM DA fluorescence in the animal tissues were imaged by a confocal microscopy. (**C, D**) Representative fluorescent image with depth color-coded MIP of the body skin labeled by DAF-FM DA in zebrafish and axolotl, respectively. The magnified MIPs of the areas within the white boxes are shown in the right panels of each image. White dotted lines indicate the orientation of collagen fibers. (**E**) Representative fluorescent images of the tendon labeled by DAF-FM (green) in the zebrafish fin bones. Fin bones were stained with Alizarin Red (magenta). (**F**) Representative fluorescent images of the tendon and ligament labeled by DAF-FM DA (green) in the axolotl forelimb digits. te, tendon; li, ligament. Scale bar = 50 μm (B, C and F) and 200 μm (E).

**Fig. S7.**
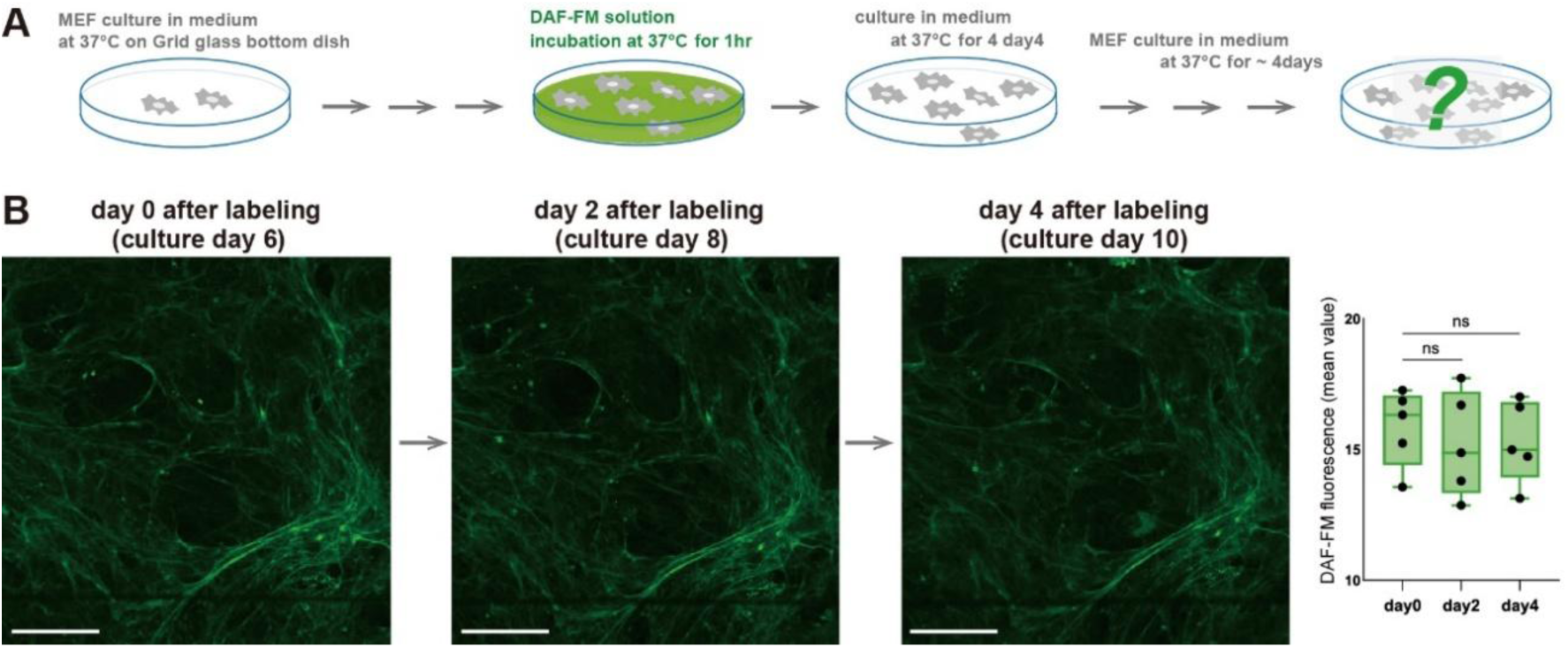
Fluorescence of the collagen fibers labeled with DAF-FM hardly fade after washout. (**A**) Schematic diagram of the temporal observation after DAF-FM staining for the collagen fibers formed by MEFs. (**B**) Representative fluorescent images of the collagen fibers labeled with DAF-FM at day 0, day 2 and day4 after labeling. Fluorescent mean intensity values of the DAF-FM at each time point are shown in the right panel. Scale bar = 100 μm.

**Fig. S8.**
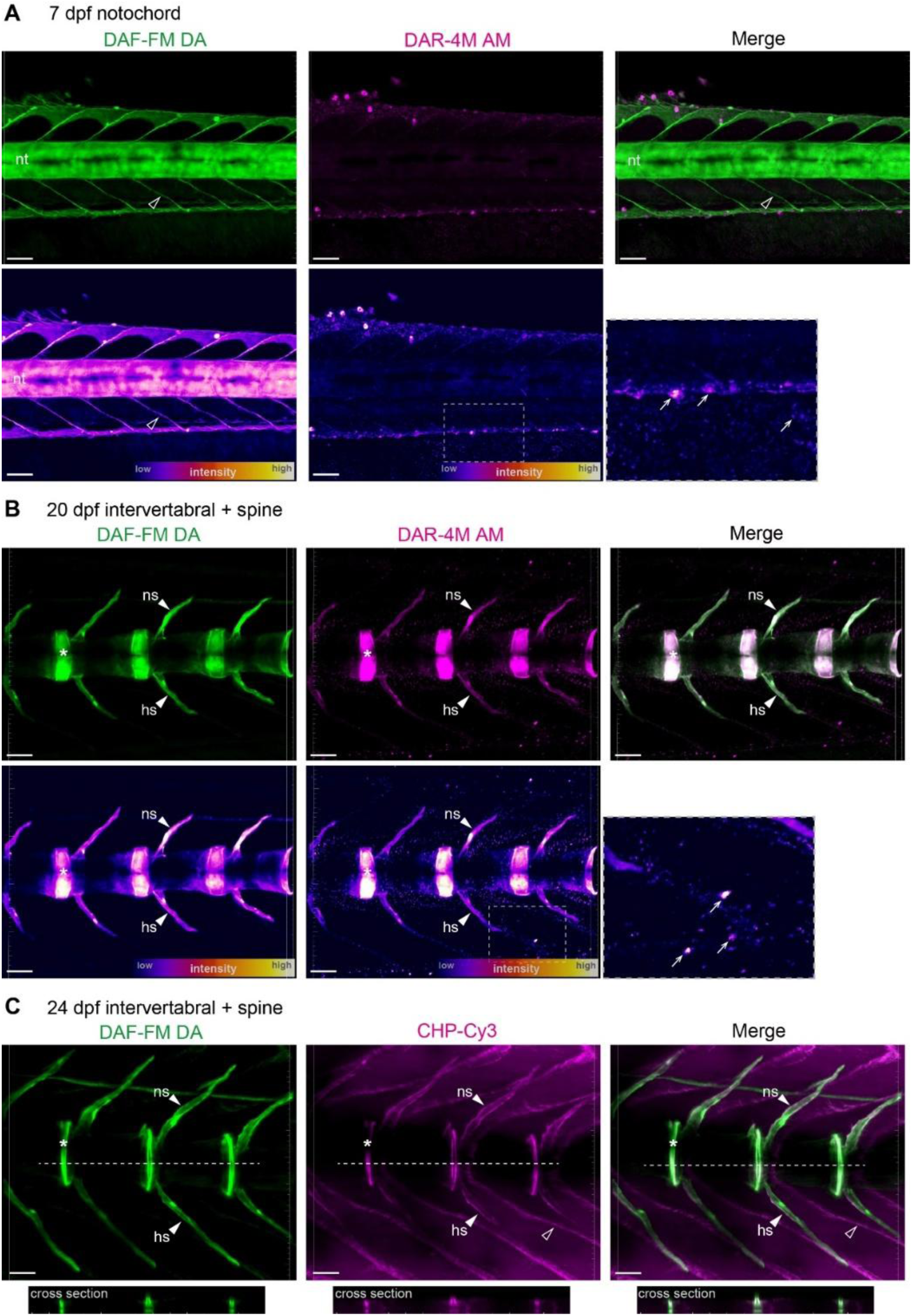
Reactivity of DAF-FM DA and DAR-4M AM with zebrafish notochord and vertebral bones. (**A**) Representative confocal images of the collagen fibers in the notochord at 7 dpf simultaneously labeled with DAF-FM DA and DAR-4M AM. Each image with fluorescence intensity color-coded MIPs is shown in the lower panels. (**B**) Representative confocal images of the collagen fibers in the intervertebral regions and spines at 20 dpf simultaneously labeled with DAF-FM DA and DAR-4M AM. Each image with fluorescence intensity color-coded MIPs is shown in the lower panels. (**C**) Representative confocal images of the collagen fibers in the intervertebral regions and spines at 24 dpf co-labeled with DAF-FM DA and CHP-Cy3. Cross section images at the position of the white dotted lines in each fluorescent image are shown in the lower panels. nt, notochord; ns, neural spine; hs, hemal spine. Open arrowheads indicate the myoseptum tendon, arrows indicate unknown nonspecific signals detected in DAR-4M AM labeled samples, and asterisks indicate the intervertebral discs. Scale bar = 50 μm.

## DISCUSSION

In this study, we established a simple and robust method for visualizing collagen fibers in cultured cells and vertebrate tissues using the small-molecule fluorescent probes DAF-FM and DAR-4M. Collagen fibers are deeply embedded within tissues and form complex three-dimensional networks, making whole-mount staining challenging. Our approach overcomes these limitations by enabling efficient penetration of fluorescent probes and stable labeling of collagen fibers without requiring specialized pretreatment. Even thick tissues such as fibrocartilage in the intervertebral disc could be clearly visualized by overnight immersion in low concentrations of DAF-FM DA (Fig. 4A-C), demonstrating the broad applicability of this method for imaging collagen-rich structures.

A central finding of this study is that DAF-FM and DAR-4M fluorescently label collagen fibers through a mechanism independent of nitric oxide. We identified that DAF-FM covalently reacts with aldehyde groups on allysine residues in the telopeptide domain of type I collagen (Fig. 2E-G), which serve as precursors for LOX-mediated cross-link formation. This annulation reaction provides a highly stable fluorescent signal that persists through tissue clearing (Fig. 4A-C), SDS-PAGE, and membrane transfer (Fig. 2D, Fig. S5B), enabling long-term visualization of collagen fibers *in vitro* and *in vivo*. These properties distinguish DAF-FM from previously reported aldehyde-reactive probes that rely on reversible hydrazone or oxime formation, and highlight the unique chemical stability of DAF-FM–mediated labeling. Several recent studies focusing on LOX-mediated cross-linking have proposed chemical approaches for detecting collagen fibers using aldehyde-reactive probes that form hydrazone or oxime linkages with allysine residues (21–25). Although these reactions enable selective detection of collagen, the resulting bonds are, in principle, reversible. In contrast, DAF-FM undergoes an annulation reaction with aldehyde groups on allysine, generating a highly stable adduct that resists elimination (Fig. 2H). Consistent with this chemical stability, fluorescence from DAF-FM–labeled collagen persisted for several days in vitro and remained detectable for days to weeks in both cultured cells and living tissues.

Our method also provides a powerful strategy for analyzing collagen fiber dynamics. Because DAF-FM selectively labels newly secreted collagen prior to cross-link maturation, fluorescence remains tightly associated with individual fibers and does not diffuse after washout. By combining DAF-FM and DAR-4M in a pulse–chase format, we were able to distinguish newly formed collagen fibers from pre-existing ones, offering a unique opportunity to monitor fiber growth and remodeling over time. This capability is difficult to achieve with approaches using fluorescent protein–tagged collagens (*16–19*, *45*, *46*), which may interfere with fibril assembly due to their large globular domains and are limited to observing collagen distribution within a restricted temporal window. In contrast, our approach labels normally secreted collagen molecules with minimal steric hindrance, enabling dynamic visualization of fiber formation in both cultured cells and living vertebrates (Fig. 5).

The chemical accessibility, low cytotoxicity, and high tissue permeability of DAF-FM and DAR-4M further enhance their utility for collagen imaging. Because fluorescence is generated only upon covalent reaction with target aldehyde groups, background signal from unreacted probes is minimal, facilitating high-contrast imaging of collagen fibers in diverse tissue environments. These features make the probes well suited for applications requiring deep-tissue visualization, long-term imaging, or quantitative analysis of collagen organization.

Together, our findings establish DAF-FM and DAR-4M as versatile tools for visualizing collagen fibers and tracking their dynamic behavior in vertebrate tissues. By targeting an intermediate structure in the collagen cross-linking pathway, this method provides a unique window into the early stages of fibril formation and enables three-dimensional imaging of collagen networks with high stability and specificity. We anticipate that this approach will facilitate future studies of collagen fiber growth during tissue development and remodeling, and will contribute to a deeper understanding of pathological conditions such as fibrosis, in which dysregulated collagen production and turnover play central roles.

### Limitations of the study

This study demonstrates that DAF-FM and DAR-4M label collagen fibers through covalent binding to allysine residues, but the detailed reaction mechanism, including reaction kinetics and potential interactions with other aldehyde-containing molecules, remains to be fully characterized. The probes do not distinguish among fibrillar collagen types, such as types I, II, III, V, and XI (4, 5, 47), and labeling efficiency may vary depending on tissue type, developmental stage, and extracellular environment, as factors including tissue permeability and collagen density could influence probe accessibility. Although the dual-probe pulse–chase strategy enables temporal visualization of newly synthesized collagen fibers, it does not provide direct quantitative measurements of collagen synthesis or turnover rates. In addition, the reactivity of DAR-4M was lower at certain developmental stages for reasons that remain unclear (Fig. S8A), and the generalizability of both probes across broader tissue contexts and additional vertebrate systems remains to be evaluated. Further studies will be required to refine the chemical specificity of these probes and to expand the range of biological applications.

## ACKNOWLEDGMENTS

We are grateful to the members of the Kondo Laboratory (Laboratory of Pattern Formation) at Osaka University and the members of the Kuroda Laboratory (Laboratory of Morphogenesis) at the JT Biohistory Research Hall. We thank Dr. Ritsuko Suyama and Dr. Ritsuko Morita for managing the confocal microscopes at the Graduate School of Frontier Biosciences, Osaka University. We acknowledge the Leica Imaging Lab and Nikon Imaging Center at Osaka University for their support in obtaining fluorescence imaging data.

This research was funded by the Japan Society for the Promotion of Science (JSPS) KAKENHI, Grant Number 21K06200, and the Japan Science and Technology Agency (JST) FOREST Program, Grant Number JPMJFR224P.

## AUTHOR CONTRIBUTIONS

Conceptualization: J.K. and T.K. Methodology: J.K., K.F., S.F., and A.H. Investigation: J.K., K.F., and Y.T. Visualization: J.K., K.F., and Y.T. Supervision: J.K. and T.K. Writing—original draft: J.K. Writing—review and editing: J.K., K. F., S. F., A.H., Y.T., and T.K. Funding acquisition: J.K.

## SUPPLEMENTAL INFORMATION

Figures S1–S8

Movie S1. Sequential confocal microscopy images of cartilage primordia and notochord located inside the tail of an E14.5 mouse embryo.

Movie S2. Sequential confocal microscopy images of fibrocartilage oriented in the intervertebral disc inside the tail of a P14 mouse.

## MATERIALS AND METHODS

### Animal maintenance and tissue preparation

Mouse tissues were prepared at Osaka Medical and Pharmaceutical University. All mouse experiments were approved by the Institutional Review Board of Osaka Medical and Pharmaceutical University and performed in accordance with the Guide for Animal Care and Use of Laboratory Animals of Osaka Medical and Pharmaceutical University. ICR mice were purchased from Japan SLC, Inc. (Shizuoka, Japan). Mice were euthanized under deep anesthesia using isoflurane inhalation for adults and hypothermia for pups and embryos. Tissues were harvested in phosphate buffered saline (PBS) and utilized for whole-mount staining or fixed with 4% paraformaldehyde in 0.1 M phosphate buffer (pH 7.4) for 2 d at 4 °C. The zebrafish were maintained under standard laboratory conditions and treated as previously described (*48*). AB strains were used as wild-type zebrafish. Albino axolotls were raised and maintained under 14 h light/10 h dark cycles at 20 °C and fed commercial granular solid food twice a day. Zebrafish and axolotls were anesthetized with tricaine (MS-222) at optimal concentrations according to their body size. All experiments involving these animals were approved by the Animal Care and Use Committee at Osaka University.

### Microscopy and image analysis

Fluorescent images were obtained using confocal microscopes LSM 780 (Carl Zeiss), STELLARIS8 (Leica), and FV1000 (Olympus), as well as a two-photon microscope (A1R MP+/Ti2-E, Nikon). 890 nm laser excitation and a 440 nm SP emission filter were used for the SHG imaging of collagen fibers. ZEN (Carl Zeiss), LAS X (Leica), FV10-ASW (Olympus), and Fiji were used as imaging software for z-projections. 3D image analysis was performed using Imaris 10.1.1 (Oxford Instruments). The fluorescence intensity and orientation angles of the collagen fibers were measured using the FIJI.

### DAF-FM/DAR-4M staining for the visualization of collagen fibers formed by cultured cells

DAF-FM (Goryo Chemical, SK1003-01) and DAR-4M (Goryo Chemical, SK1005-01) were used to stain collagen fibers produced by mouse embryonic fibroblasts (MEF) (Reprocell, RCHEFC003). The cells were cultured in Dulbecco’s Modified Eagle’s Medium (DMEM, FUJIFILM Wako Pure Chemical Corporation) supplemented with 10% fetal bovine serum (FBS), 200 μM L-ascorbic acid phosphate magnesium salt n-hydrate (FUJIFILM Wako Pure Chemical Corporation), 100 U/mL penicillin, and 100 µg/mL streptomycin (Sigma-Aldrich) at 37 °C in a 5% CO_2_ atmosphere. 35 mm glass bottom dishes coated with 0.1% gelatin solution (Nacalai Tesque, 19895-75) were used for the MEF culture experiment. For the staining of collagen fibers deposited around the cultured cells, DAF-FM and DAR-4M were diluted with culture medium (DMEM with 10 % FBS) and adjusted to concentrations of 6.9 μM and 9.8 μM, respectively. Before staining with DAF-FM or DAR-4M, the cells were washed three times with DMEM without l-ascorbic acid phosphate magnesium salt n-hydrate. The cells were then incubated with the diluted DAF-FM or DAR-4M solutions for 1 h at 37 °C. After incubation, samples stained with probes were washed with fresh medium and observed under a confocal microscope. For pulse-chase observation of collagen fibers formed *in vitro* culture conditions, the cells were first incubated with DAF-FM solution, then after 4 days of culture with fresh medium, they were next incubated with DAR-4M solution for 4 days. For confocal microscopy analysis, the fluorescent signals of the collagen fibers stained with DAF-FM and DAR-4M were detected using 488 nm and 561 nm lasers, respectively.

### DAF-FM staining of the purified collagen

DAF-FM (Goryo Chemical, SK1003-01) was used to stain purified collagen substrates. Gelatin (Nacalai Tesque, 19895-75), collagen type I (Nitta Gelatin, Cellmatrix Type I-P) and collagen type ǁ (AteloCell, CL-22) were prepared on 35 mm glass bottom dishes as substrates for DAF-FM staining. Gelatin and collagen type ǁ were adjusted to concentrations of 0.1% and 0.03 mg/ml, respectively, and coated on glass. Collagen type I was adjusted to a concentration of 1.8 mg/ml and neutralized with 1N NaOH to form a gel on glass.

Each substrate was incubated with the DAF-FM solution diluted with DMEM with 10% FBS, adjusted to a concentration of 6.9 μM, for 1 h at 37 °C. After incubation, samples stained with DAF-FM were washed with fresh medium and observed under a confocal microscope.

### DAF-FM DA/DAR-4M AM staining for the visualization of collagen fibers in vertebrate tissues

DAF-FM DA (Goryo Chemical, SK1004-01) and DAR-4M AM (Goryo Chemical, SK1006-01) were used for whole-mount staining to visualize vertebrate tissues. For the staining of whole-mount mouse tissues, DAF-FM DA was diluted with PBS, adjusted to concentrations of 5 μM, and used for staining. Mouse embryos or dissected tissues, including P0–P1 skins and P14 tails, were incubated in the staining solution under dark conditions and treated for 12 h at room temperature. After the staining, they were washed with PBS and fixed with 4% paraformaldehyde (PFA) in PBS O/N at 4 °C. After fixation, the cells were treated with RapiClear 1.52 (SJL, RC152001) O/N at room temperature for optical tissue clearing and were observed using a confocal or two-photon microscope. For the staining of whole-mount juvenile of axolotls, DAF-FM DA and DAR-4M AM were diluted with breeding water, adjusted to concentrations of 5 μM and 10 μM, respectively, and used for staining. Living juvenile of axolotls were bathed in the staining solution under dark conditions and treated for 12 h at 20 °C. After the staining, they were anesthetized with tricaine (MS-222) at an optimal concentration and fixed with 4% PFA in PBS O/N at 4 °C. After fixation, the skin was dissected and observed under a confocal microscope. Axolotl forelimbs were observed under a two-photon microscope after subsequent transparency treatment with RapiClear 1.49 (SJL, RC149001). For the staining of whole-mount zebrafish larvae, DAF-FM DA and DAR-4M AM were diluted with breeding water, adjusted to concentrations of 5 μM and 10 μM, respectively, and used for staining. Living zebrafish larvae were bathed in DAF-FM DA solution in the dark and treated for 12 h at room temperature. After the staining, they were anesthetized with MS-222 at an optimal concentration and fixed with 4% PFA in PBS O/N at 4 °C. After fixation, tissues, including skin, tendons, and bones, were observed using a confocal microscope. For the pulse-chase observation of vertebral collagen fibers, zebrafish larvae at 7dpf were first stained with DAF-FM DA, then after 2 weeks of breeding in a circulating tank, they were next stained with DAR-4M AM. For observation under a confocal microscope, the fluorescent signals of the collagen fibers stained with DAF-FM DA and DAR-4M AM were detected using 488 nm and 561 nm lasers, respectively.

### Detection of DAF-FM modification of lysine at the telopeptide domains of type I collagen

The cell/matrix layers were sequentially digested with bacterial collagenase and pepsin to analyze the DAF-FM modifications of telopeptidyl lysine in type I collagen, as previously reported (*49*). In brief, the samples were heated at 80 °C for 30 min, and digestion with 0.01 mg/mL of recombinant collagenase from *Grimontia hollisae* (Nippi, Tokyo, Japan) (*50*) was performed in 100 mM Tris-HCl/5 mM CaCl_2_ (pH 7.5) at 37 °C for 16 h. After addition of acetic acid (final 0.5 M), digestion with 0.01 mg/mL of pepsin was further performed at 37 °C for 16 h. The peptide solutions were subjected to LC-MS analysis on a maXis II quadrupole time-of-flight mass spectrometer (Bruker Daltonics, Bremen, Germany) coupled to a Shimadzu Prominence UFLC-XR system (Shimadzu, Kyoto, Japan) using an Ascentis Express C18 HPLC column (5 µm particle size, L × I.D. 150 mm × 2.1 mm; Supelco, Bellefonte, PA, USA) (*49*). Peaks of peptides containing lysine or allysine labeled with DAF-FM (+392.061 Da) were detected in the extracted ion chromatograms.

### Antibody staining of collagen

For the antibody staining for collagen type I, the cultured MEFs were fixed with 4% PFA in PBS and blocked with 1% bovine serum albumin (BSA)/PBS. After blocking, they were incubated with anti-mouse collagen type I rabbit polyclonal antibody (Rockland Inc, 600-401-103-0.1, 1:100 dilution) solution in 1% BSA/PBS O/N at 4 °C. Next day, they were washed with PBS and incubated with goat anti-rabbit IgG (H+L) antibody, FITC conjugate (Invitrogen, 65-6111, 1:200 dilution) solution in 1% BSA/PBS for 2 h at room temperature. For the antibody staining of collagen type ǁ, the mouse E14.5 embryos were fixed with 4% PFA in PBS. After fixation, the tails were dissected and sequentially immersed in 10%, 20%, and 30% sucrose solutions in PBS, and in a 1:1 solution of 30% sucrose and Tissue-Tek O.C.T Compound (Sakura Finetek Japan, 4583). Subsequently, they were embedded in O.C.T compound and frozen on dry ice. Tissues were sectioned at a thickness of 10 µm using a cryomicrotome CM1850 (Leica). The cryosectioned samples were blocked with 2% bovine serum albumin/PBS (BSA)/PBS. After blocking, they were incubated with anti-chick collagen type ǁ mouse monoclonal antibody (DSHB, ǁ-ǁ6B3, 1:100 dilution) solution in 2% BSA/PBS O/N at 4 °C. Next day, they were washed with PBS and incubated with goat anti-mouse IgG (H+L) antibody, Alexa 594 conjugate (Invitrogen, A11020, 1:200 dilution) solution in 1% BSA/PBS for 1 h at room temperature. Immunofluorescence images were obtained by confocal microscopy.

### Staining with denatured collagen-binding peptide

For staining of the collagen fibers deposited around cultured cells, the cells were treated with phosphate-buffered saline (PBS) heated at 95 °C for 1 min to heat-denature extracellular collagen. They were then fixed with a 4% PFA and blocked with 2% BSA/PBS. After blocking, they were incubated with 5 µg/mL of Bin*d*COL, biotin-conjugated (Funakoshi, FDV-0035) in 1% BSA/PBS or 3 µg/mL of fluorescein-conjugated soCMP6-7(Glu)2 in 1% BSA/PBS O/N at 4 °C and washed with PBS (*35*). The cell samples incubated with Bin*d*COL solution were stained with streptavidin, Alexa 647 conjugate (Invitrogen, S32357, 1:200 dilution) solution in 1% BSA/PBS for 1 h at room temperature and washed with PBS after staining. Finally, the fluorescence signals of the denatured collagen fibers were imaged by confocal microscopy. To staining of the collagen fibers distributed in zebrafish tissues, the tissues were fixed with a 4% PFA. After fixation, the cells were heat-treated in a thermal bath at 80 °C for 10 min to denature the extracellular collagen. They were then blocked with 2% BSA/PBS, incubated with 5 µM of CHP-Cy3 in 2% BSA/PBS O/N at 4 °C, and then washed with PBS. Finally, the fluorescence signals of the denatured collagen fibers were imaged by confocal microscopy.

### Staining of collagen deposited around BAPN-treated cultured cells

Previously established MEF clones were used for staining (*51*). The cells were cultured in DMEM with 10% FBS and 200 μM L-ascorbic acid phosphate magnesium salt n-hydrate at 37 °C in a 5% CO_2_ atmosphere. The medium was replaced with HFDM-1(+) (Cell Science & Technology Institute Inc.) containing 100 U/mL penicillin and 100 µg/mL streptomycin after the cells had reached confluence in 35 mm glass-bottom dishes. Confluent MEFs were incubated in HFDM-1(+) with or without the addition of 500 µM 3-aminopropionitrile fumarate (BAPN, Sigma-Aldrich) for 2 d in 35 mm glass-bottom dishes. Subsequently, the cells were washed with PBS and incubated in a medium containing DAF-FM/DMSO (final concentration of 6.9 µM DAF-FM, and 0.1% DMSO) or DMSO (final concentration of 0.1%) at a dilution of 1/1000 for an additional hour. In cases where BAPN was added, it was introduced to a concentration of 500 µM. After washing the cells with PBS, they were fixed in a 4% paraformaldehyde phosphate buffer solution for 10 min. The cells were then washed with PBS and observed under a confocal laser microscope.

### Staining of cellular actin and nucleus

For staining the actin cytoskeleton and cell nuclei of the cultured MEFs, the cells were fixed with 4% PFA. After fixation, the cells were washed with PBS and incubated with a solution of Phalloidin-fluor 594 conjugated (AAT Bioquest; 1:300 dilution) and Hoechst (Dojindo; 1:500 dilution) in PBS for 2 h at room temperature. For staining of cell nuclei in frozen sections of mouse tail tissues, the samples were fixed with 4% PFA. After fixation, they were washed with PBS and incubated with a solution of Hoechst (Dojindo; 1:500 dilution) in PBS O/N at 4 °C. For staining of cell nuclei in P14 the tail and P1 mouse skin, the tissue samples were fixed with 4% PFA. After fixation, they were washed with PBS and incubated with a solution of Syto 82 (Invitrogen; 1:1000 dilution) in PBS O/N at 4 °C. Each sample was washed with PBS after staining, and nuclear fluorescence was captured using confocal microscopy.

### BrdU incorporation assay

MEFs were incubated in culture medium for 6 d and then incubated in DAF-FM solution at a concentration of 6.9 μM for 1 h. After staining with DAF-FM, cells were washed with fresh medium and treated for 12 h in medium containing BrdU (Abcam, ab142567) adjusted to a concentration of 10 μM. Cells were fixed with cold 70% EtOH for 5 min at room temperature, and then treated with 0.1N HCl solution containing 0.1% Triton for 30 min at 37 °C to increase permeability of the cell nuclei. After several washes in PBS, cell samples were blocked for 1 h with a 1% BSA/PBS solution for antibody staining. Mouse monoclonal anti-BrdU antibody (Molecular Probe, A21300, 1:200 dilution) was used as the primary antibody, and cell samples were incubated O/N at 4 °C in the antibody solution containing 1% BSA/PBS. The next day, they were washed with PBS and incubated with goat anti-mouse IgG antibody, Alexa 594 conjugated (Invitrogen, A11020, 1:200 dilution) solution with 1% BSA/PBS for 2 h at room temperature. Cell nuclei were simultaneously stained with Hoechst (Dojindo; 1:500 dilution), and the percentage of BrdU-positive nuclei was compared between cells without (control) and with DAF-FM staining.

### Cell viability assay

MEFs were incubated in culture medium for 7 days and then incubated with DAF-FM solution at a concentration of 6.9 μM for 1 h. After staining with DAF-FM, cells were washed in fresh medium and incubated in Syto 82 (Invitrogen, S11363, 1:1000 dilution) solution at 37 °C for 30 min to label the nuclei of living cells. Cells were then washed in fresh medium and incubated in Nuclear Blue (AAT Bioquest, Live or Dead Cell Viability Assay Kit, 22788, 1:200 dilution) solution for 30 minutes at 37 °C to label the nuclei of dead cells. After nuclear staining, the cells were washed with PBS and the percentage of Nuclear Blue-positive nuclei was compared between cells without (control) and with DAF-FM staining.

### Drug treatment for NO removal and NOS inhibition

MEFs were cultured for 10 days in the culture medium, as described above. They were then treated under the following three different conditions: DMSO (0.1% in DMEM) as control, 2-(4-Carboxyphenyl)-4,4,5,5-tetramethylimidazoline-1-oxyl-3-oxide (C-PTIO, Dojindo) (500 μM in DMEM) known as a NO remover (*52*), and N^G^-nitro-L-arginine methyl ester hydrochloride (L-NAME, Dojindo) (500 μM in DMEM) known as an inhibitor of NOS (*53*). In the control and L-NAME treatment experiment, after 24 h of treatment, MEFs were incubated for 1 h at 37 °C in DAF-FM solution (6.9 μM) diluted with DMEM containing the corresponding drug (0.1% DMSO and 500 μM L-NAME). In C-PTIO treatment experiment, after 30 min of the treatment, MEFs were incubated in DAF-FM solution (6.9 μM) diluted with DMEM containing C-PTIO (500 μM) for 1 h at 37 °C. After DAF-FM staining, fluorescent images of the collagen fibers produced by MEFs were obtained using a confocal microscope, and the fluorescence intensity values were measured using FIJI image analysis software.

### Western blot analysis of collagen in cell layers

As in the experiment for staining collagen deposited around BAPN-treated cultured cells, confluent MEFs in 35 mm dishes were incubated in HFDM-1(+) with or without BAPN for 2 days and then treated with DAF-FM/DMSO or DMSO for 1 h. Following a wash with PBS, the cell layers were dissolved in SDS-PAGE sample buffer (50 mM Tris-HCl [pH 6.7], 10% glycerol, and 2% SDS) and heated at 95 °C for 5 minutes. The protein concentration of these SDS samples was determined using Pierce BCA protein assay kit (Thermo Fisher Scientific, Waltham, MA, USA). SDS-PAGE was performed on a 5% polyacrylamide gel with 91 mM 1,4-dithiothreitol (DTT)-reduced or non-reduced samples, and proteins on the gel were transferred onto nitrocellulose membranes. Fluorescent bands were visualized using a CCD imager LAS-3000 (Fujifilm, Tokyo, Japan). Subsequently, the membranes were blocked with 5% skim milk/Tris-buffered saline (TBS; 50 mM Tris-HCl pH 7.4, 150 mM NaCl) and washed with TBS. They were treated with 1 µg/mL of biotin-conjugated Bin*d*COL in 2% skim milk/TBS to detect collagen polypeptides (*35*). The membranes were washed with TBS, treated with streptavidin-HRP (Thermo Fisher Scientific, 1:5000 dilution) in 2% skim milk/TBS, and washed with TBS containing 0.1% Tween-20. Collagen bands were detected with a CCD imager LAS-3000 using Pierce ECL western blotting substrate kit (Thermo Fisher Scientific).

### SDS-PAGE analysis of cell layers treated with pepsin

Confluent MEFs in 35 mm dishes were incubated in HFDM-1(+) containing DAF-FM/DMSO (final concentration 6.9 µM DAF-FM, 0.1% DMSO) or DMSO (final concentration 0.1% DMSO) for 2 d. The cell layers were washed with PBS and those collected with cell scrapers were treated with 0.1 M HCl, with or without 100 µg/mL pepsin (Sigma-Aldrich), at 4 °C for 16 h. After neutralization with NaOH, the samples were mixed with 5 × SDS sample buffer and heated at 95 °C for 5 min. Proteins in 91 mM DTT-reduced or non-reduced samples were separated by SDS-PAGE on an 8% polyacrylamide gel, and fluorescent bands were visualized using a CCD imager LAS-3000. Protein bands were visualized using Coomassie Brilliant Blue R-250 staining.

### Quantification and Statistical analysis

Statistical analyses were performed using GraphPad Prism version 9.5.1 (731). All data are presented as mean ± SD. An unpaired two-tailed Student’s *t*-test was used to assess the statistical significance of differences between the means of two independent groups. Analysis of variance (ANOVA) followed by Tukey’s multiple comparison test were conducted to evaluate the statistical significance of the differences among the three groups.

